# Exploring Environmental Coverages of Species: A New Variable Selection Methodology for Rulesets from the Genetic Algorithm for Ruleset Prediction

**DOI:** 10.1101/531079

**Authors:** Anni Yang, Juan Pablo Gomez, Jason K. Blackburn

## Abstract

Variable selection for, and determination of variable importance within, species distribution models (SDMs) remain an important area of research with continuing challenges. Most SDM algorithms provide normally exhaustive searches through variable space, however, selecting variables to include in models is a first challenge. The estimation of the explanatory power of variables and the selection of the most appropriate variable set within models can be a second challenge. Although some SDMs incorporate the variable selection rubric inside the algorithms, there is no integrated rubric to evaluate the variable importance in the Genetic Algorithm for Ruleset Production (GARP). Here, we designed a novel variable selection methodology based on the rulesets generated from a GARP experiment. The importance of the variables in a GARP experiment can be estimated based on the consideration of the prevalence of each environmental variable in the dominant presence rules of the best subset of models and its coverage. We tested the performance of this variable selection method based on simulated species with both weak and strong responses to simulated environmental covariates. The variable selection method generally performed well during the simulations with over 2/3 of the trials correctly identifying most covariates. We then predict the distribution of *Bacillus anthracis* (the bacterium that causes anthrax) in the continental United States (US) and apply our variable selection procedure as a real-world example. We found that the distribution of *B. anthracis* was primarily determined by organic content, soil pH, calcic vertisols, vegetation, sand fraction, elevation, and seasonality in temperature and moisture.

## 1. Introduction

Species distribution models (SDMs; i.e. ecological niche models [ENMs]) have been widely applied in ecology, biogeography, conservation biology, evolution, and epidemiology over the past several decades (Larson et al., 2010; Ostfeld et al., 2005; Pearson and Dawson, 2003; Peterson and Vieglais, 2001). Modeling a species’ geographic distribution relies on some form of pattern-recognition based on non-random association between the geographic occurrences of a species and environmental conditions that support its survival under the ecological niche theory (Araujo and Guisan, 2006; Hutchinson, 1957). The ecological niche of a species can be defined as the environmental conditions that allow the population to be maintained without immigration (Grinnell, 1917; Pulliam, 1988) and can be described by an n-dimensional hyper-volume of environmental covariates that determine the ecological space of the species (Hutchinson, 1957). Hence, the accuracy of predicted distributions is primarily driven by the adequacy of environmental covariates used in the models (Araujo and Guisan, 2006; Austin, 2007). Species’ distributions and their environmental requirements can be veiled or misleading due to the selection of inappropriate predictors (Araujo and Guisan, 2006). Incorporating the suitable covariates in ecological niche modeling experiments remains an important area of research with continuing challenges.

Most SDM algorithms use exhaustive searches through variable space (in multiple combinations) in order to identify the variables that define a species’ distribution. As the most biologically-based decision in SDMs, the selection of environmental covariates should primarily depend on the knowledge of the adaption of species’ physiology to the ecological or biological conditions (ecophysiological or biophysiological processes) that govern the relationships between a species and the environment (Austin, 2007). However, this information is difficult to obtain in many cases, especially for some poorly understood species. With a large number of potential predictors, including biotic and abiotic, direct and indirect factors, which influence species’ responses to environmental gradients and available resources (Austin and Van Niel, 2011), some crucial questions arise, like “how many variables are enough” and “which variables need to be included” (Araujo and Guisan, 2006; Huston, 2002). The evaluation of variable contributions within SDMs is an alternative to quantify the relationship between the species survival and environment to understand the ecological requirements of a species. The estimation of variable contribution in the SDMs provides an objective metric to infer the strength of species response to the environmental conditions, which can help to hypothesize about the ecophysiological processes determining the geographical distributions and understand some basic biology of the species (Araujo and Guisan, 2006). Finally, the variables contributing most are selected to interpret the species’ ecological niche and predict the most likely distribution (species range).

The estimation of each variable’s explanatory power and the selection of the optimal variable set within models, however, can be challenging for some species distribution modelling approaches, such as the Genetic Algorithm for Ruleset Production (GARP). GARP is a common technique for predicting species distributions based on presence-only data via an algorithm employing a superset of logistic regression, range and negated range rules, and atomic (bioclim) rules (Stockwell, 1999). GARP experiments can employ the Jackknife procedure (Levine et al., 2007; Peterson and Cohoon, 1999; Thomasson and Blouin-Demers, 2015), but there is no easy way and rubric for the estimation of variable contribution. Levine et al. (2009) presented a method for performing a statistically based comparison between the comprehensive map (i.e. N variables) and jackknifed maps (i.e. N-1variables) generated from GARP to determine the optimal ecological parameters for predicting human monkeypox disease. The larger differences found between the output from an experiment with all models and the map produced from a jackknifed experiment, the greater the contribution the reduced variable made in those experiments (Levine et al., 2009). However, this estimation relies on the prediction performance of GARP and assumes that the comprehensive map, as the base map, represents the geographic distribution predicted by the “true” fundamental niche. Also, the computational intensity for massive iterations of the jackknife procedure makes variable selection difficult when there is a large set of potential environmental covariates. Alternatively, Sweeney et al. (2007) employed an external classification and regression tree (CART) to select the optimal environmental layers to be used in GARP experiments to model the distribution of *Anopheles punctulatus* in Australia. However, GARP and CART use different algorithms to determine relationships between species’ occurrences and environmental covariates. GARP includes logistic regression and range envelopes, while CART constructs decision trees by making binary splits of the covariates. These differences in algorithms may result in different estimations of variable explanatory power and therefore the variable set selected by CART may not be optimal for GARP.

Exploring the variable space that defines the ecological niche of a species can help us in understanding the underlying ecophysiological processes of the species’ distribution. Here, we present a novel variable selection methodology for GARP based on the exploration of the GARP rulesets to consider the explanatory power of variables within a modeling experiment and the biological information within the experiment using those variables. We base our variable selection process mainly in two metrics: 1) the prevalence of each environmental variable in the dominant presence rules of the best model subset from a GARP experiment, and 2) the variables’ median range in those rules. In this study, we explain in detail the new variable selection procedures and test its performance using simulations and provide a real-world case study for exploring ecological requirements and predicting the distributions of the *Bacillus anthracis* in the continental US using a bioclimatic variable set recently introduced to the modeling community.

## 2. Materials and Methods

### 2.1. GARP

GARP is a presence-only iterative modeling algorithm that searches for non-random relationships between point occurrence data and environmental covariates. For this study, we use DesktopGARP (DG) version 1.1.3 to perform GARP experiments. The procedure for running a GARP experiment is demonstrated in Fig. 1. Initially, we split the occurrence data into external training and testing sets. The external training set is inputted in DG for model building, while the testing set is withheld for external model accuracy tests to evaluate the performance of GARP experiment. Each properly executed GARP experiment will include multiple models and each will have a ruleset with 50 rules predicting presence or absence (note: there are GARP implementations in openModeller allowing the user to control the number of rules). There are four types of rules (range, negated range, atomic, or logit) described as the if/then logic statements. Range rules specify the envelope with upper and lower bounds for the presence of the species (e.g. IF temperature = [10.2 – 13.5°C] AND NDVI = [0.15 – 0.23] THEN species = PRESENCE). Negated range rules define the conditions outside of variable ranges (e.g. IF NOT temperature = [10.2 – 13.5°C] AND NDVI = [0.15 – 0.23] THEN species = ABSENCE). Logit rules employ logistic regression to determine the relationship between the species occurrence and covariates (e.g. IF temperature*0.0037 + NDVI*0.57 THEN species = PRESENCE). The presence or absence of the species in the logit rule type is determined based on the probability of the occurrence of the species predicted by the logistic regression with the threshold of 0.5.

**Fig. 1.**
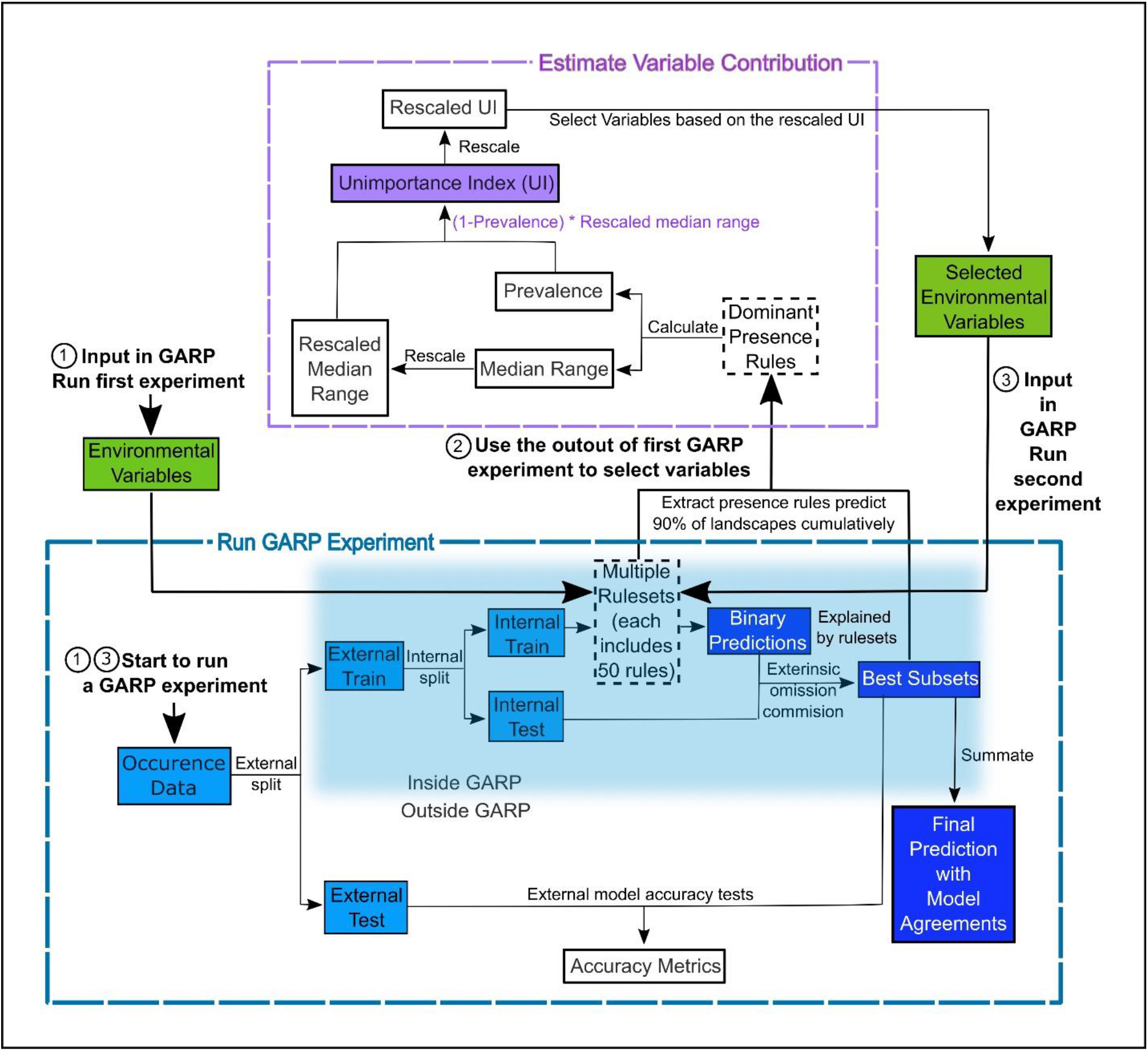
Flowchart depicting the procedure to run a GARP experiment and estimate variable contribution. There are three steps for predicting species distribution and selecting variables selected via Unimportance Index (UI). First, run a complete GARP experiment with the full variable set. Second, use the output of the first GARP experiment to rank and select variables based on UI. Third, input the important variables in GARP to run the second GARP experiment to predict the species distributions.

Atomic rules use specific values of the covariates to determine the presence of the species (e.g. IF temperature = 12.5°C AND NDVI = 0.19 THEN species = PRESENCE). Those rules are developed and tested internally using random draws of presence points from the known occurrences and random draws of the background space representing absences (i.e. pseudo-absences). An internal chi-square test built on the predicted and observed values is used to evaluate the quality of each rule at predicting presence or absence with the user’s pre-defined proportion of input data (internal testing set). GARP can accept, modify or delete rules using deletions, insertions, cross-overs, among other types of mutations to improve predictive accuracy in a genetic fashion.

Once a ruleset is developed, it is projected onto the geography of the study area to develop a presence/absence map describing the species’ potential geographic distribution, e.g. Blackburn (2006), Joyner (2010), and Stockwell (1999). Given the iterative nature of GARP, the model does not arrive at a single solution. DG splits input occurrence data into training and testing sets inside the software for model evaluation and incorporates a “best subset” procedure, which would select the best subset of models based on two criteria: omission (false negative) and commission (false positive; percent of pixels predicted present) rates. Such calculations are performed on each individual model and the “best subset” procedure selects a user defined number of models based on specific omission and commission values. Here, experiments were setup to run up to 200 models, we selected 20 models with no more than 10% “extrinsic” omission rate, which is calculated from the internal testing set. A median commission percentage is then calculated for the 20 low-omission models. Investigators can define the percentage (defaulted to 50%; 10 models) of the low-omission models that have individual commission closest to the median to be selected as the best subset (McNyset and Blackburn, 2006). Finally, the best subset with 10 best presence-absence predictions can be summed and mapped on the landscape with model agreements indicating the likelihood of the species presences. GARP has been shown to perform well across the spectrum of species’ prevalence on the landscape from rare to common making it useful for management oriented studies focused on relating geographic potential to management or conservation needs (Peterson et al., 2007). A more extensive description of GARP’s modeling framework and test of its performance can be found elsewhere (Anderson et al., 2003; Martinez-Meyer et al., 2006; Peterson and Cohoon, 1999; Stockwell, 1999), and in this study, we limit our objectives to describe the variable selection procedure.

### 2.2. Conceptual Framework for variable selection procedures

We designed a new variable selection methodology to estimate variable contributions to species distributions in GARP. We used accuracy metrics (omission and commission rates and area under the curve (AUC)) to select the best subset of models (rulesets) in the GARP experiment. We measured the variable contributions based on two criteria: 1) the prevalence of the variable in the dominant presence rules and 2) the scaled median range for those variables across the rules within the best subset of the GARP experiment.

The prevalence of a variable in the dominant presence rules of the best subset is defined as the frequency with which the variable predicts the presence of the species in the dominant presence rules of the best subset (See Equ. 1). With the best subset process activated, DG selects a set of best models as described above. The dominant presence rules in the best subset are defined as a subset of rules that cumulatively predict the over 90% of the species’ presence on the landscape in the top-selected 10-model subset (Mullins et al., 2011). Those rules represent the primary suitable environmental conditions that define the core of the ecological niche of the species (based on the set of variables available) but does not take into account rare situations in which species are occasionally or temporarily present. Here we only analyzed presence rules, since absence rules tend to have wide median ranges. We defined prevalence as:

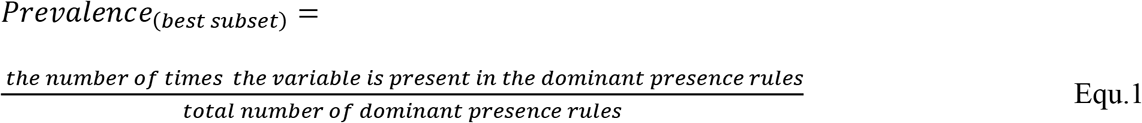

The high prevalence rate of a variable indicates that the variable is frequently used to predict the presence of the species in the best subset. Thus, a variable with a higher prevalence rate suggests the variable is relatively more important in the GARP experiment.

The median range of a variable is defined as the difference between the median values from a set of maximum and minimum values of this variable in the dominant presence rules from the best subset (Joyner, 2010). For different types of rules, the maximum and minimum values are extracted in different ways. In range and negated range rules, the maximum and minimum values are extracted directly from the upper and lower boundaries recorded in the rulesets. For the logit rules, the maximum and minimum values are extracted from the landscape where those logit rules are used to predict the presence of the species via zonal statistics. For atomic rules, the specific values of the covariates that predict the presence of the species are directly extracted from the rules. We then compare the extracted value of the atomic rules with the maximum and minimum values from other types of rules to evaluate whether it fell inside the coverage. To quantitatively compare the median ranges of different variables, we scale the median range of each variable from 0 to 1 (Barro et al., 2016). A variable with a wide median range indicates that the presence of species is not sensitive to this predictor, while a variable with a narrow median range suggests that the occurrence of the species is constrained to specific conditions regarding the covariate (Barro et al., 2016; Mullins et al., 2011).

We measured the variable contribution to GARP based on an Unimportance Index (UI) to consider both criteria, the prevalence rate and scaled median range. The UI of each covariate is calculated as the multiplication of the scaled median range and the probability that the variable is not used to predict the presence of the species in the dominant presence rules of the best subset (Equ. 2). This multiplication would help to combine and balance both criteria. Variables with less contribution to a GARP experiment are defined as the ones with wider median range and lower prevalence. Therefore, the larger the UI value is, the less contribution the associated variable brings to the model. To clearly compare and evaluate variable contribution we finally rescaled the UI to 0-1 following Equ. 3:

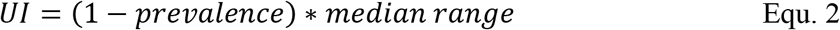

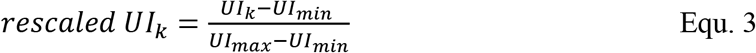

where UI_k_ is the unimportance index for covariate k; UI_max_ and UI_min_ are the maximum and minimum value of the UIs for the covariates in the variable set, respectively. This procedure of the estimation of variable contributions are shown in Fig. 1 and programmed in “GARPTools” R-package (available at https://github.com/cghaase/GARPTools).

### 2.3. Testing the performance of the new variable selection procedure using simulations

#### 2.3.1. Simulating the species and sampling it

To test the performance of the aforementioned variable selection method we first generated ten normally distributed environmental covariates with spatial autocorrelation on a 10.5 * 10.5 degree landscape at a 0.01 degree resolution (Fig. A. 1). Five of those covariates were simulated using an exponential variogram model with a range of 10, sill of 1, and nugget of 0, the others used a spherical variogram model with a range of 6, sill of 1, and nugget of 0. Next, we simulated 200 species using three variables from the entire set drawn at random without replacement. The probability of occurrence was computed as:

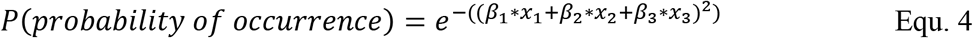

where β_1_, β_2_, and β_3_ are the coefficient that determines the influence of each covariate on the species distribution and x_1_, x_2_, and x_3_ are the environmental covariates. The three selected variables used in species distribution simulation were recorded for further validation of the performance of the variable selection procedures. Once we obtained the probability surface on the landscape, we used it as the success probability of a Bernoulli random trial to obtain the true distribution (Elith and Leathwick, 2009). The three coefficients for each species were sampled from a normal distribution under two scenarios. The first represents a scenario in which the environmental covariates weakly define the species distribution. In this case, we sampled the coefficients from a normal distribution with mean of one and standard deviation of 0.5. For the second scenario we assumed that the coefficients had a stronger effect on the distribution of the species such that the coefficients were normally distributed with mean of five and a standard deviation of 0.5. We simulated 100 species using the weak effect coefficients and 100 using the strong effect. Finally, we randomly extracted 50 presence locations from the centroid of the grid cells of the realized distribution for each species as the presence-only data to input in GARP.

#### 2.3.2. Testing the variable selection performance

To test the performance of the UI, we used the full set of ten environmental variables and the 50 presence points sampled from the species distribution to generate a GARP experiment for each species. Here, since the true distributions of the simulated species is known, we can directly compare the predictions with true distributions without withholding part of data for external model validation. We set the training/testing data split to 75%/25% inside DG. To maximize GARP performance, model runs were set to a maximum of 1,000 iterations or until convergence of 0.01. The best subset procedure selected ten best models under a 10% extrinsic omission threshold and a 50% commission threshold (Fielding and Bell, 1997). Those 10-model best subsets were added together using GARPTools R-package.

For each of the 200 species we calculated the UI for all the ten variables used in model development and recorded the three variables with the lowest UI (i.e. the three variables with highest contribution to the predicted distributions). We evaluated the performance of the model and the UI by counting the number of variables *r* (*r* = 0,1,2,3) correctly identified by the model for each of the species. Next we counted the number of species *s* (*s* = 0, 1, 2,…, *S*) with *r* = 0, 1, 2, and 3. Finally, we compared the distribution of s to the distribution generated by drawing three variables at random out of the ten used to generate each SDM. The probability of *r* = 0, 1, 2, 3 is given by

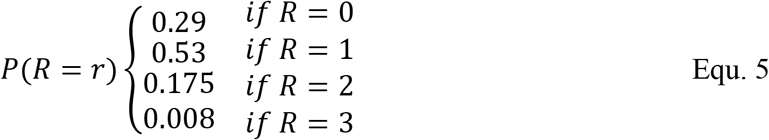

We then used a one tailed Pearson’s chi-squared statistic to compare the expected and observed number of cases with zero, one, two, and three variables being correctly identified for all the 200 simulated species and for each weak and strong effect scenario separately (see Appendix B for proof of how probabilities were derived).

### 2.4. Case study: modeling *Bacillus anthracis* in the continental US

Applications of SDMs to pathogens or disease systems remain an important tool for estimating disease distributions or mapping risk areas. Understanding variable contribution can assist on evaluating biological information within models and how those compare to real-world knowledge of pathogen or host/vector biology. To explicitly demonstrate the use of the new variable selection procedure, we provide a real-world case study for exploring the ecological requirements and distributions of the *B. anthracis* in the continental US.

Anthrax, a zoonotic disease, primarily affects wildlife and livestock and secondarily afflicts humans nearly worldwide (Alexander et al.,2012). *Bacillus anthracis*, the causative agent of anthrax, is a spore-forming bacterium, which is endemic to specific soil environments and can persist for extended periods of time (years to decades) (Van Ness, 1971). Several ecological niche modeling studies have defined the ecological niche as a narrow range of moderate NDVI (indicative of grasslands) with limited annual precipitation and high soil pH (Barro et al., 2016; Blackburn et al., 2007; Joyner, 2010; Mullins et al., 2011). Anthrax is an established disease in the US (Stein, 1945) and still remains endemic in some parts of the country, such as the recent outbreaks in Montana in 2008 and 2010 (Blackburn et al., 2014a; Morris et al., 2016) and the enzootic zone of West Texas (Blackburn et al., 2014b).

#### 2.4.1. Data

We adopted the historical anthrax outbreak data (305 cases) from Blackburn et al. (2007). The outbreaks in eastern Oklahoma were excluded from this study, since the environmental conditions in that region are not suitable for the survival of *B. anthracis* spores, and those occurrence of the outbreaks and temporary suitable environment were suggested to result from anthropogenic activities (Blackburn et al., 2007; Van Ness, 1959). We used 26 climatic and biophysical covariates as the environmental coverages for modelling distribution of *B. anthracis*. The details of data and sources are shown in Table 1. All environmental layers were resampled to 2.5 arcminute resolution. Given the resolution of the environmental layers, the 305 anthrax outbreak cases represented 175 unique pixel cells which were selected using the spatially unique routine in GARPTools.

**Table 1.** Environmental variables used for *B. anthracis* GARP experiment.

| Environmental Layer (unit) | Names | Resolution | Source |
| --- | --- | --- | --- |
| Elevation (meter) | Alt | 1 km | WorldClim<sup>a</sup> |
| Bioclimatic data (°C or kg of water/<br>kg of air) | Bio 1-19 | 2.5 arcminute | MERRAclim<sup>b</sup> |
| Mean NDVI (no unit) | wd0114a0 | 1 km | TALA<sup>c</sup> |
| NDVI annual amplitude (no unit) | wd0114a1 | 1 km | TALA |
| Top soil pH (no unit) | pH | 1 km | SoilGrids<sup>d</sup> |
| Sand fraction in top soil (% weight) | sand fraction | 1 km | SoilGrids |
| Calcium Vertisols (% weight) | calcium vertisol | 1 km | SoilGrids |
| Top soil organic content (g per kg) | organic content | 1 km | SoilGrids |

*Note: a) the WorldClim elevation data were accessed from worldclim.org/ (Hijmans et al., 2005); b) the MERRAclim dataset from the 2000s decade with the mean humidity version was downloaded from https://data.dryad.org/ (Vega et al., 2016; 2017); c) NDVI measurements were accessed from the Trypanosomiasis and Land Use in Africa (TALA) research group (Oxford, United Kingdom; Hay et al., 2006); d) Four soil layers were obtained from SoilGrids website (https://soilgrids.org/). All the data were accessed on Sep 21, 2018.*

#### 2.4.2. Variable selection based on UI to predict *Bacillus anthracis*

To explore the environmental coverages for *B. anthracis*, we followed a similar procedure as for the simulated species. We first input all 26 environmental covariates in GARP. Since the true distribution of the species is unknown, and to validate the predicted distributions from GARP, we split the 175 spatially unique anthrax occurrence data into external training/testing set with 75%/25% ratio prior to model construction (Fig. 1). We built the GARP model following the parameterization in Blackburn et al. (2007). In a first GARP experiment, we calculated the UI for each of the 26 variables and assumed them to be important if the UI value was smaller than 0.5. Finally, we re-ran the GARP experiment using only the variables identified to be important.

Predictive accuracy for the best subsets from the GARP experiment with the UI-based reduced variable set was evaluated using a combination of AUC, omission, and commission rates based on the external testing dataset (Lim and Klein, 2006; Peterson et al., 2007). The AUC, although not an ideal metric for accuracy estimation (Lobo et al., 2008), is useful to identify models that perform well (Hanley and McNeil, 1982; Mullins et al., 2013; Sloyer et al., 2018). The 10-model best subset from the UI-based experiment was summated to map the potential geographic distribution of *B. anthracis* for the continental US.

## 3. Results

### 3.1. Simulated species and variable selection performance in simulation scenarios

Examples for the probability maps of species distributions, binary occurrence maps simulated with weak and strong correlations, and GARP predictions based on those simulated species are illustrated in Fig. 2. We found that UI and GARP performed well during the simulations. For the 200 simulated species we found that the observed number of species with *r* = 0, 1, 2, 3 does not follow the distribution of random draws (*χ*^2^ = 724.3, n = 200, df = 3, p < 0.0001) and in particular the observed number of species with *r* =2 and *r* = 3 is significantly higher than expected by chance (Table 2). We found a similar result when analyzing separately the species in which environmental covariates were assumed to have a weak and strong effect on the geographic distribution (Table 2; weak: *χ*^2^ = 367.2, n = 100, df = 3, p < 0.0001; strong: *χ*^2^ = 360.1, n = 100, df = 3, p < 0.0001). Finally, we found no differences in the observed number of species with *r* = 0, 1, 2, 3, when comparing the species simulated using strong and weak coefficients (*χ*^2^ = 2.64, df = 3, p = 0.45).

**Fig. 2.**
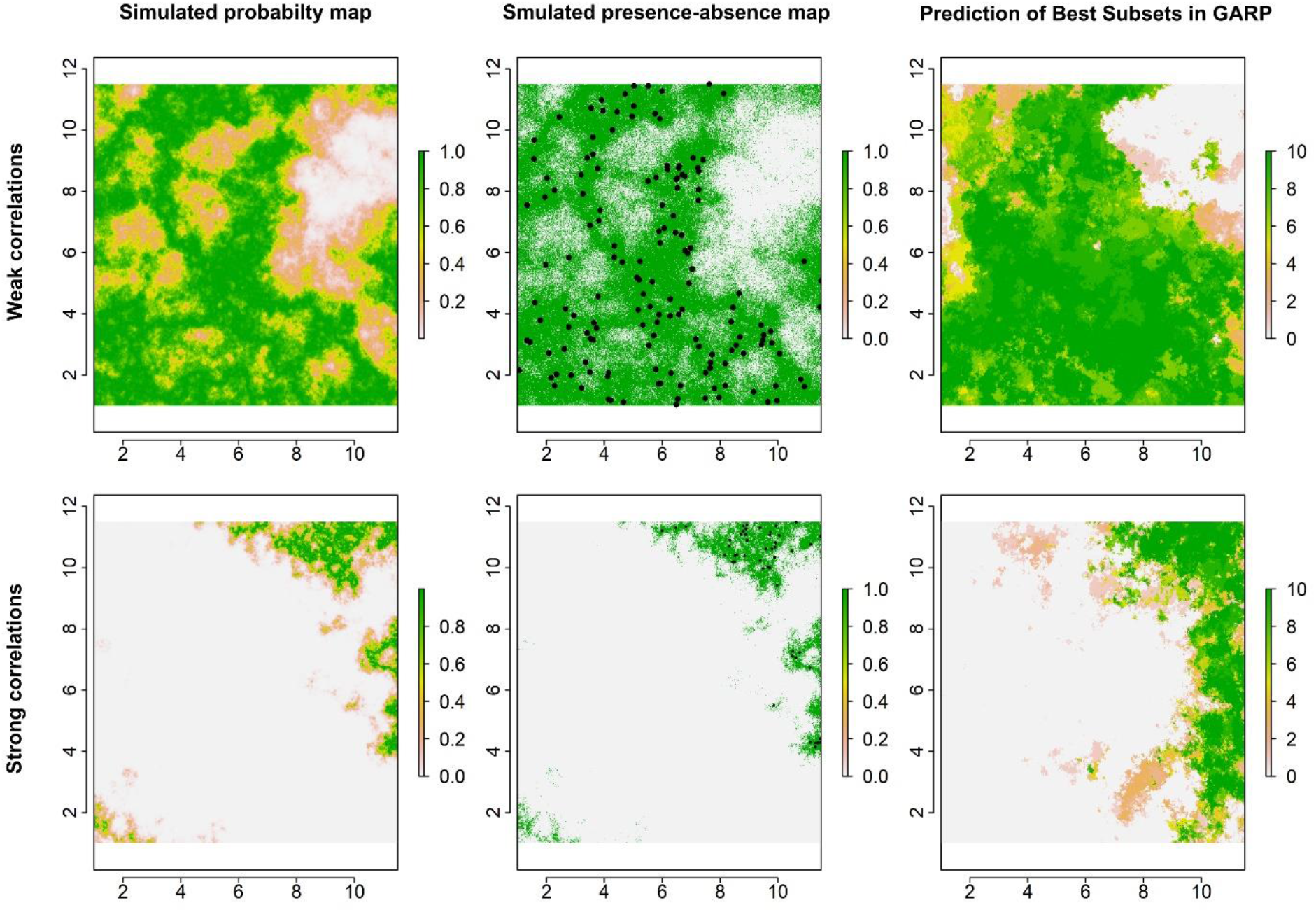
Simulated species distributions, occurrence (presence-absence) maps, and GARP prediction map for the best subset under the two scenarios where the correlation between species occurrence and environment are weak and strong; the black points are the presence locations extracted from occurrence map for modelling species distributions in GARP.

**Table 2.**
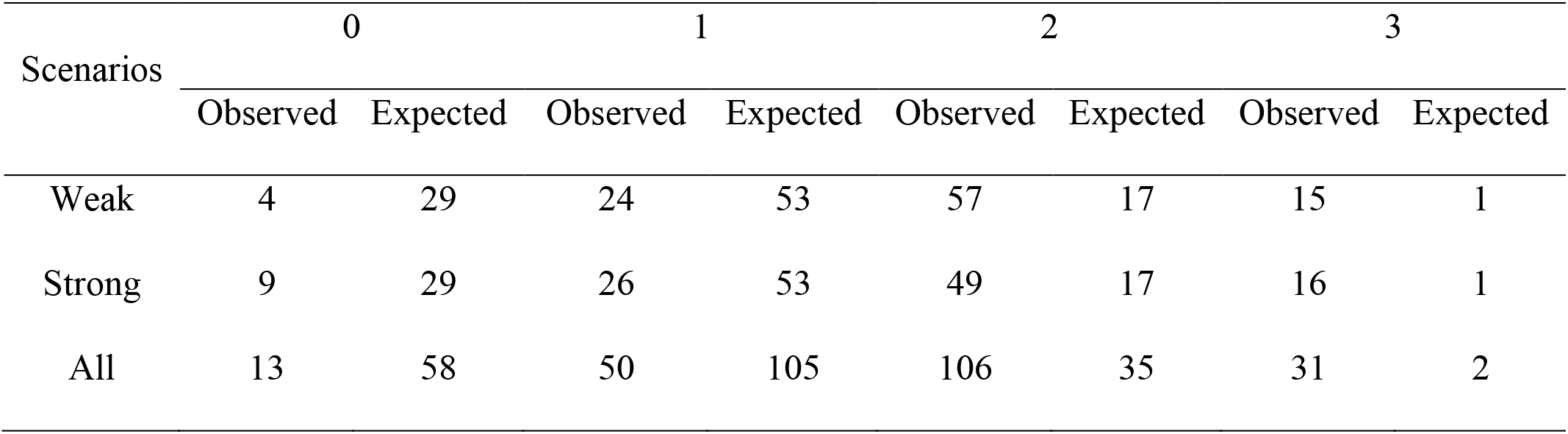
Summary of the observed and expected number of species for which the variable selection method correctly identified zero, one, two or three out of three variables used to simulate the species distribution. The counts are tallied for 200 simulated species (All) and separated by the 100 species for which we selected Weak and Strong influence of the environmental variables on determining the species distribution.

### 3.2. Ecological requirements and distributions of *B. anthracis*

We selected 12 variables with UI less than 0.5, including the climatic (temperature and moisture) seasonality, elevation, mean NDVI, seasonality of NDVI, organic contents, calcic vertisols, pH, and sand fractions (Table 3). AUC value of the GARP experiment with the reduced variable set was 0.86 (Table 4). The total and average omission rates of this best subset were 0.02% and 5.11%, respectively, and the total and average commission rates were 21.55% and 10.14%, respectively (Table 4).

**Table 3.**
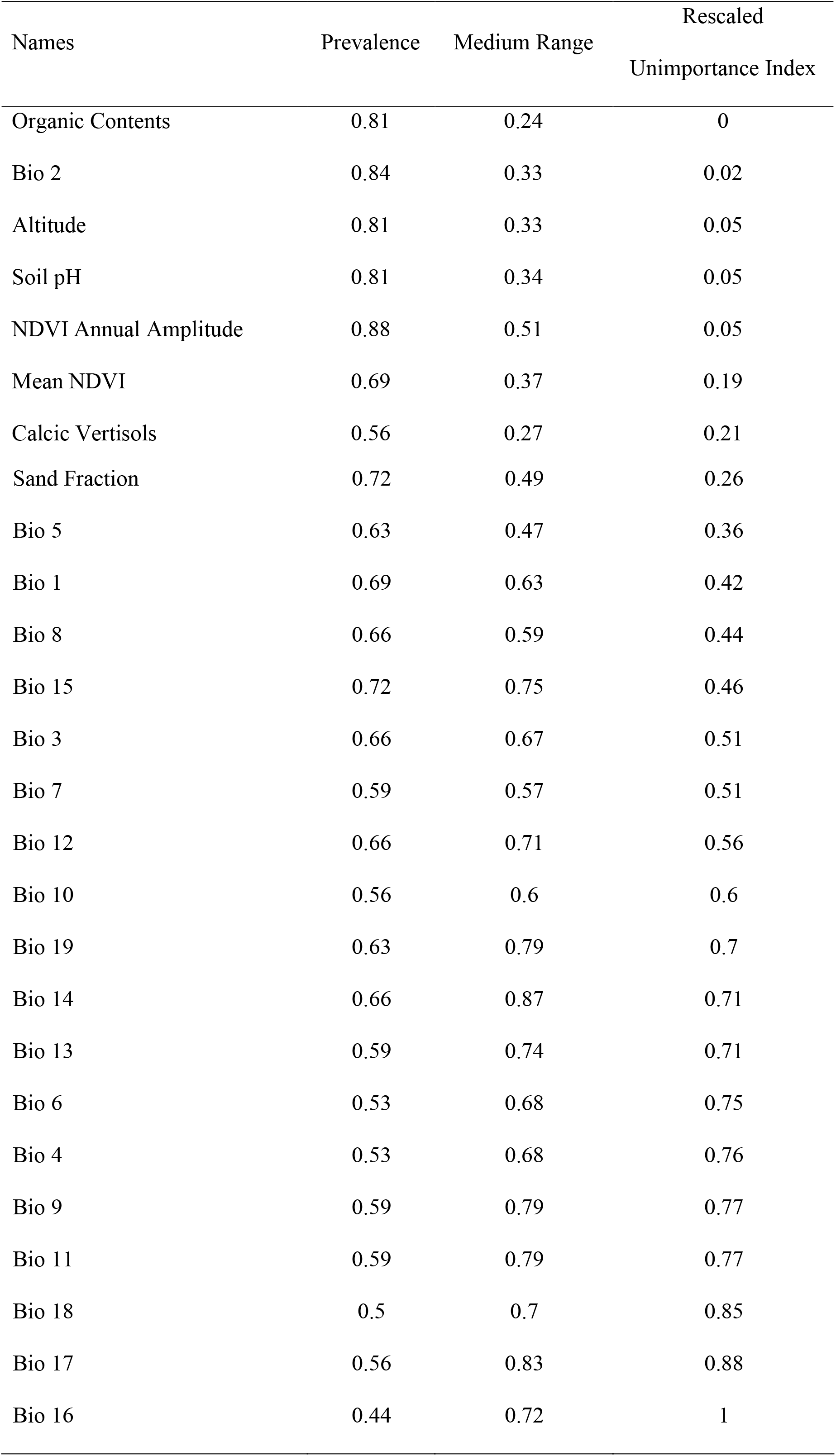
Estimation of variable contribution for the *B. anthracis* in GARP experiment.

**Table 4.** Accuracy metrics for the *B. anthracis* GARP species distribution model.

| Metric | Model Specifications |
| --- | --- |
| Num. of points in external training set | 132 |
| Num. of points in external testing set | 43 |
| Total omission | 0.02% |
| Average omission | 5.11% |
| Total commission | 21.55% |
| Average commission | 10.14% |
| AUC | 0.86 |

The GARP experiment with the reduced variable set predicted presence of *B. anthracis* primarily along a north-south corridor starting from the Dakotas, eastern Montana, and western Minnesota southward through western Wyoming, western Nebraska, eastern Colorado, western Kansas, eastern Oklahoma, and into the New Mexico and western Texas (Fig. 3). The north-south corridor also expands westward into western Washington and Oregon through southern Idaho. The distribution was predicted in some patches of Nevada, Utah, Arizona, and southwestern California. There were also some small areas along the shorelines of the Great Lakes in eastern Wisconsin, eastern Michigan, and northwestern Ohio and northeastern Indiana. Fig. 4 illustrates the scaled median ranges and coverages of variables in the dominant presence rules of the best subset in GARP model with the reduced variable set. The variable with the narrowest range was organic content, while Bio 15 had the widest range. Calcic vertisols (0 – 7.23%), altitude (134.8 – 1321.95 m), mean NDVI (−0.1 – 0.46) and soil pH (6.52 – 8.19) also had relatively small median ranges.

**Fig. 3.**
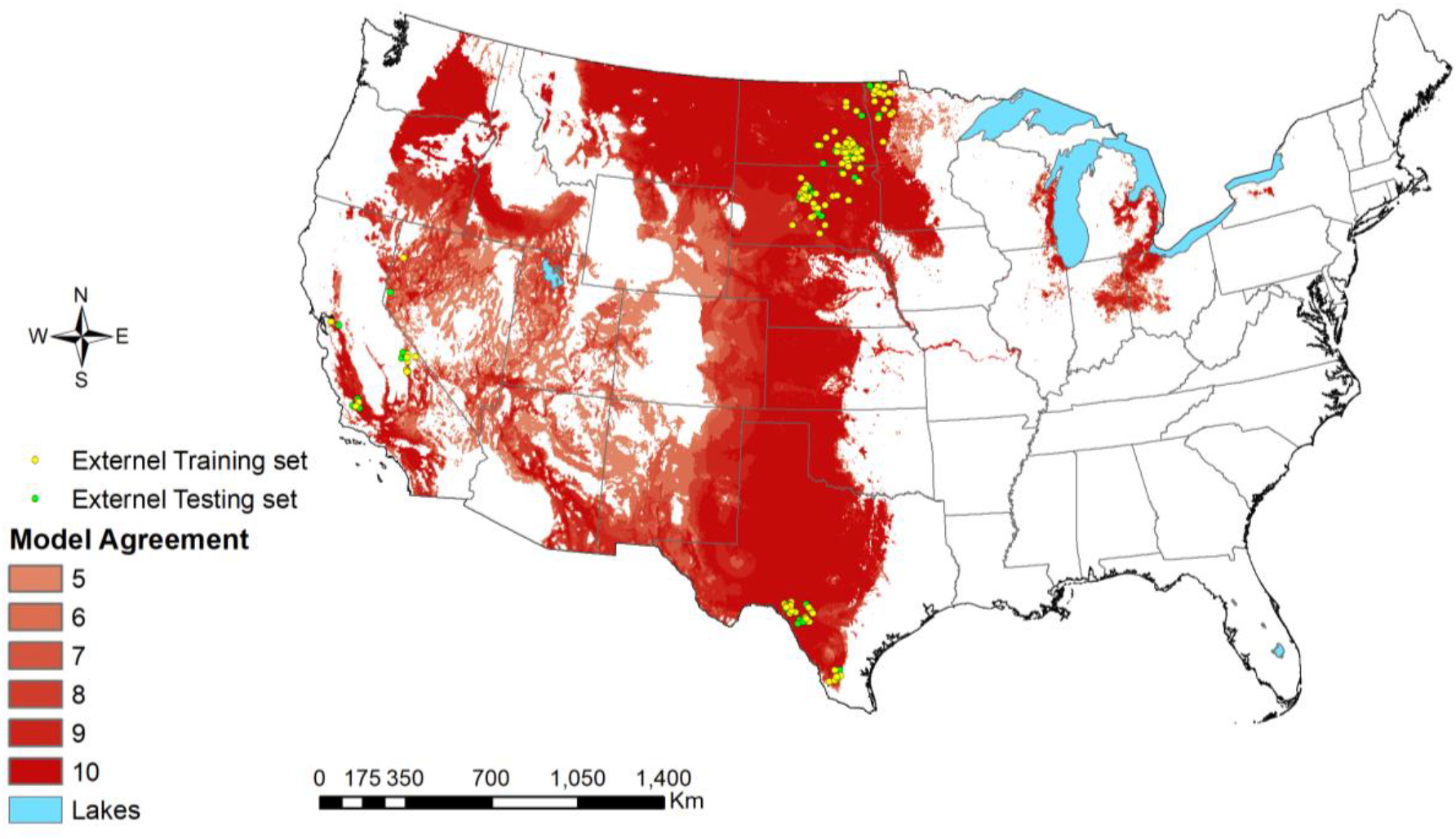
Prediction of *B. anthracis* in the continental US from the best subset in the GARP experiment using the selected variable set.

**Fig. 4.**
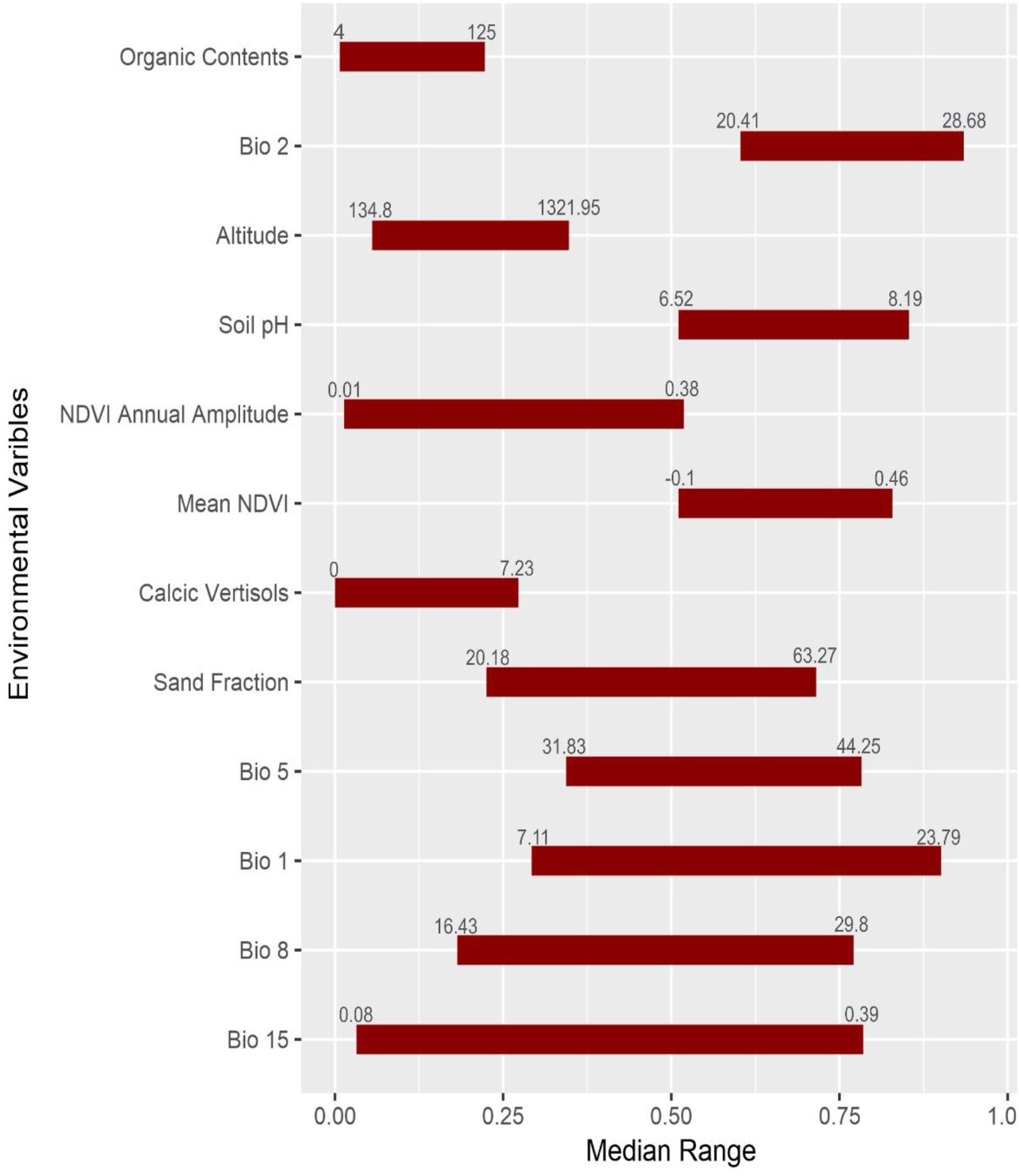
Scaled median range of the covariates from the best subset in the GARP experiment using the selected variable set; The numbers at both sides of the bar represent the real value of the upper and lower bound of coverage.

## 4. Discussion

In this study, we present a new variable selection rubric for GARP based on prevalence rates and median ranges of the variables in the dominant presence rules in best subsets. Overall, the variable selection methodology performed well by identifying the important ecological variables defining the distribution of the simulated species. We found a high probability of identifying all or most of the variables that are important to the distributions of those species, irrespective of the relative influence of the variables on determining the distribution. In over 65% of the cases, our UI correctly identified at least two of the three variables defining the species environmental envelope. In the real-world case study, we identified that 12 of 26 were of high importance in determining the distribution of the *B. anthracis* in the US. The important variables included temperature and moisture seasonality, some soil conditions, and vegetation index.

Our new methodology for estimating variable contribution in GARP was developed considering the explanatory power within a modeling experiment measured by the frequencies the variables are used and the biological information within the experiment using those variables. The explanatory power of the variables here were first measured by the number of times that the variables were selected to predict the presence of the species in the best subsets. This idea follows from the estimation of variable contributions in some machine learning algorithms, such as Boosted Regression Trees (BRTs) and random forests, which calculate the variable contributions based on the number of times the variable is used to split the trees (Friedman and Meulman, 2003). Additionally, the biological information within the GARP experiment was quantified by the median ranges of the variables. Variables with a narrow range of values that will predict the presence of the species suggest species distributions are sensitive to those conditions (Mullins et al., 2011). Those variables might have a higher explanatory power as they may restrict the species distribution in both ecological and geographical space. If a species has a wide tolerance to a specific variable, then this variable may necessarily have low explanatory power at least in the geographic area considered. Variables that are identified with less contributions to the model could also be important conditions for the species survival but allow a species to be widespread or are not the common requirement across the population of occurrences. UI considering both the frequency the variable used to predict species presence and biological information would help identify common conditions confining a species’ distribution, which could be used to infer the underlying biological mechanisms of species survival.

We tested the performance of the proposed variable contribution estimation method in simulated species with both weak and strong correlations between species occurrence and environmental covariates and found overall good performance. Our generation of the simulated species, although is simpler than reality, follows an ecologically realistic scenario in which species distributions are a function of multiple factors and respond to the environment under a bell curve determined by these covariates and is not limited to one type of species (Elith and Leathwick, 2009). The test of the performance of UI in different simulation scenarios evaluates its general ability of correctly identifying the primary covariates that contribute to species distributions. We found that majority of the cases in both simulation scenarios selected most (2/3 or all three) variables correctly, which indicates that our variable selection method performs well regardless of the strength of the environment in determining the species distribution. Overall, the good performance of UI indicates that this method allows the identification of the environmental variables that are important in defining a species distribution, and thus can allow us to make inferences about the physiological tolerances of the species and the dispersal abilities across a landscape.

The incorporation of the optimal variables in the model is important for making inferences about the ecology and the mechanisms determining species distributions. Including the optimal set of variables in the SDMs could increase the model accuracy and provide a better understanding of the ecological requirements for species survival. Also, filtering the most useful variables among a series of candidate variables might help to reduce noise in the predictions. In the real-world case study, we selected organic contents, calcic vertisols, sand fraction, soil pH, vegetation trend and amplitude, elevation, and trend and seasonality of temperature and moisture, to describe the ecological niche of *B. anthracis*. This selection is in line with the optimal environmental variables of the survival of *B. anthracis*, including the trend of climate, elevation, vegetation indexes, soil moisture, and pH, summarized by Hugh-Jones and Blackburn (2009). The high AUC (0.86) of GARP outputs for *B. anthracis* indicated a good performance of the model with the selected optimal variable set. Additionally, the ecological requirements of *B. anthracis* survival identified in this study support the results reported by alternative research (Blackburn et al., 2007; Hugh-Jones and Blackburn, 2009; Hugh-Jones and De Vos, 2002; Van Ness, 1971). Anthrax is known as a hot season disease (Blackburn and Goodin, 2013) and our results suggest that the spores of *B. anthracis* were found in the places with annual mean temperature ranging from 7.11 – 23.79 °C, mean diurnal ranges varying 20.41 – 28.68 °C, and the maximum temperature in the warmest quarter from 31.83 – 44.25 °C. The UI selected all soils variables and vegetation index and suggested that *B. anthracis* was predicted to be found in areas with high soil pH (6.52-8.19), low calcic vertisols (0 – 7.23%), sand fraction of 20.18 – 63.27%, organic contents ranging between 4 – 125 g/kg soil, mean vegetation index from −0.1 – 0.46, vegetation annual amplitude ranging from 0.01 – 0.38. In line with our results, high concentrations of spores have been found in black steppe soils with alkaline pH (e.g. over 6.0 recorded in Van Ness (1971); 5.5 – 7 in Kracalik et al. (2017) in Ghana), moderate in organic matter and calcium content (Hugh-Jones and Blackburn, 2009). The optimal vegetation greenness for anthrax occurrence is suggested as a narrow range of moderate NDVI (approximately 0.2 to 0.5; indicative of grasslands), e.g. 0.1 – 0.3 in Kracalik et al. (2017), 0.17 – 0.56 in Blackburn (2006).

The distribution of *B. anthracis* predicted here with the reduced variable set was similar to the predictions in Blackburn et al. (2007), except that the southern part of the corridor in this study was slightly more widespread than the previous results. Also, more areas around the Great Lakes region were predicted to be highly preferred by *B. anthracis* in our model. Those differences in the predictions might result from the different variable set and data sources used in the SDMs. Blackburn et al. (2007) used annual trend of climatic data (i.e. mean annual temperature and precipitation from the Bioclim dataset; (Hijmans et al., 2005)), elevation, mean NDVI, and soil moisture, and pH to develop the model, while this study included the seasonality in temperature and moisture from MERRAclim dataset (Vega et al., 2017), elevation, mean NDVI, seasonality of NDVI, organic contents, pH, calcic vertisols, and sand fractions based on our estimation of variable contributions. Additionally, different spatial scales can also influence the predictions. Given the modifiable areal unit problem in quantitative ecological studies (Openshaw and Taylor 1979), the values of pixels could vary with the changes of the pixel sizes. While Blackburn et al. (2007) predicted the distribution with ~8 * 8 km^2^ spatial resolution, we used a ~4.5 * 4.5 km^2^ pixel size. Despite these differences, the accuracy metrics were high and the prediction plausible.

## 5. Conclusions

The method described herein presents a procedure of evaluating variable contributions based on median range and the frequency the variable used to predict the presence of the species. This variable contribution estimation procedure was employed using GARP system, but the idea of the consideration of both the explanatory power and environmental coverage when selecting variable is highlighted and is applicable to other SDMs. The new variable selection method was tested via simulations which we found to be accurate in the identification of the important environmental variables in determining the distribution of simulated species. We employed this method to understand the ecological requirements and geographic distributions of *B. anthracis*. The optimal ecological coverages selected by the variable selection method include the seasonality of temperature and moisture, elevation, mean and seasonality of NDVI, organic contents, calcic vertisols, pH, and sand fractions. The predicted distributions were primarily restricted to central and western US. The variable selection idea presented here provides an objective way to identify the variables that are most important for predicting species distributions with GARP, which is analogous to the variable selection methods integrated in other SDM algorithms (e.g. Maxent or BRTs) and fills the gap in the practical application in the estimation of variable contributions and variable selections in GARP.

## Acknowledgements

This study was partially supported by the National Institutes of Health [grant number 1R01GM117617-01] to JKB.

## Appendices

## Appendix A. Figure for the simulated environmental layers

**Fig. A.1.**
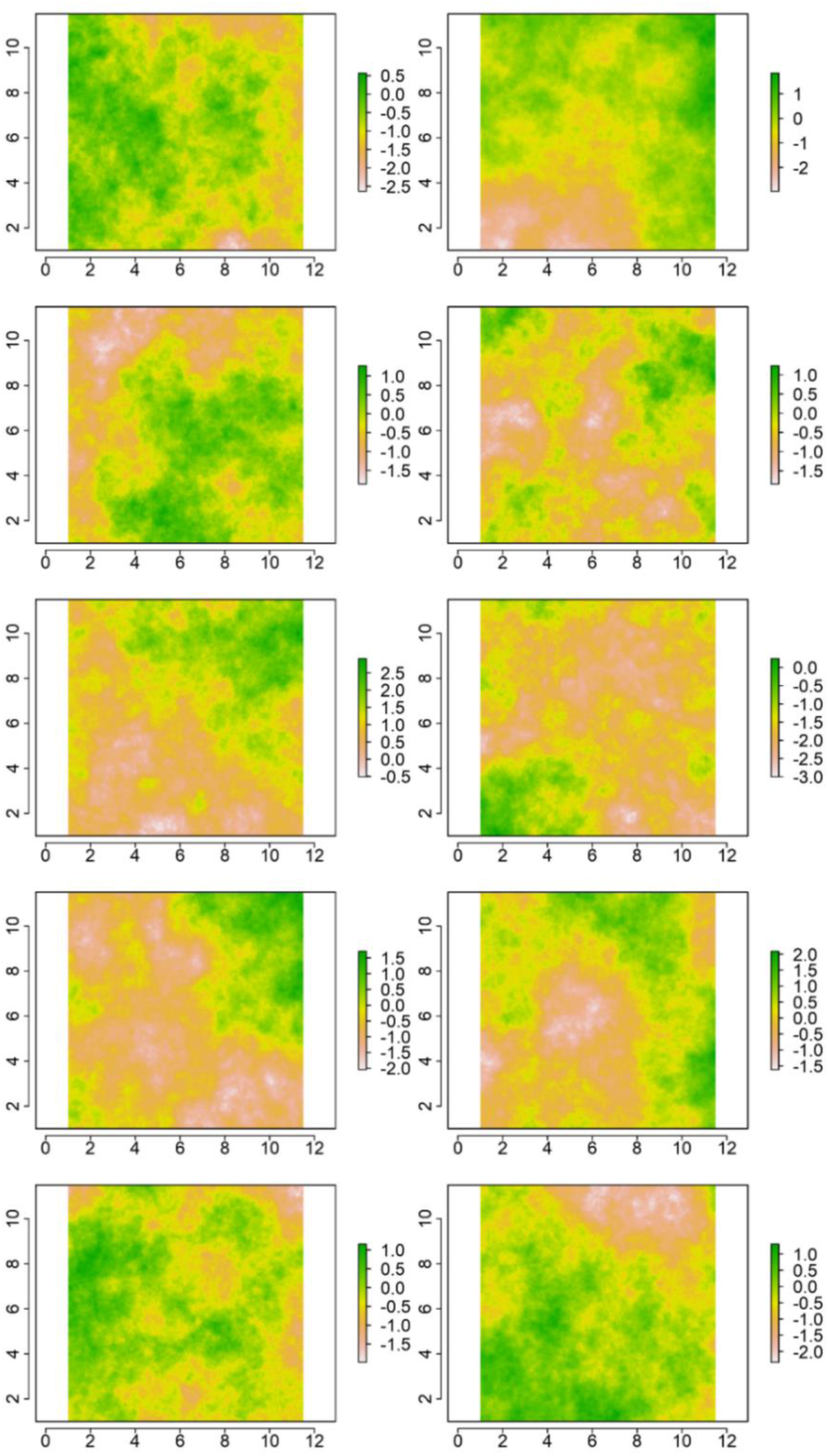
Simulated environmental layers with an extent of 10.5 * 10.5 degree and 0.01 *0.01 degree resolution; the origins of both x and y coordinates start from 1.

## Appendix B. Derivation of the probabilities *r* = 0, 1, 2, 3 based on a random draw

For *r* = 0, the probability is given by the joint probability of obtaining an incorrect variable in each of the three draws. In the first draw, there are 7 out of 10 variables that were not used to generate the species distribution thus, the probability of choosing an incorrect variable in the first draw is 7/10. Then, in the second draw, there are only 9 variables left to choose from and only 6 of them are incorrect such that the probability of obtaining an incorrect variable in the second draw is 6/9. Using the same rationale, the probability of choosing an incorrect variable in the third draw is 5/8. Thus, the probability of picking 3 incorrect variables out of 10 possible ones without replacement is just the multiplication of 7/10, 6/9 and 5/8. Thus *P*(*r* = 0) = (7/10) * (6/9’) * (5/8) ≈ 0.3. Now, for *r* = 3, using the same rationale as for *r* = 1, for the first draw there are three correct variables out of ten, in the second draw five that we chose a correct variable in the first draw there are only two out of nine left and in the third draw, given that we chose correctly the variables in the first and second draws there is only one correct variable out of eight to choose from. Thus *P*(*r* = 3) = (3/10) * (2/9) * (1/8) ≈ 0.008. For the cases in *r* = 1 and *r* =2 we need to take into account the order in which we can draw one or two correct variables. For example, for *r* = 1, we can choose the correct variable in the first, second or third draw. This means that we have three ways of choosing one variable out of ten. It is, that in the first draw we choose the correct variable and in the other two are incorrect or that we choose an incorrect variable in the first draw, the correct one in the second and an incorrect one in the third again or that we choose two incorrect variables in the first two draws and a correct one in the third draw. Let C be the draw of a correct variable and I be the draw of an incorrect variable. Thus, the chances of getting exactly one correct variable out of ten in three draws is represented by, CII, ICI, IIC. This is, *P*(*r* = 1) = (3/10) * (6/9) * (5/8) + (7/10) * (3/9) * (6/8) + (7/10) * (6/9) * (3/8) = 0.525. Similarly, for *r* =2, we have that the ways of picking two correct variables out of ten are, CCI, CIC, ICC. *P*(*r* = 2) = (3/10) * (2/9) * (7/8) + (3/10) * (7/9) * (2/8) + (7/10) * (3/9) * (2/8) = 0.175. Since the random variable *R* can only take values of 0, 1, 2, 3, the sum of the probabilities must add up to one. *P*(*P* = *r*) = *P*(*r* = 0) + *P*(*r* = 1) + *P*(*r* = 2) + *P*(*r* = 3) = 0.291 + 0.525 + 0.175 + 0.008 ≈ 1. Because of the precision with which we are defining the probabilities, the latter does not add up to one but taking into account all decimal places it does.

## Abbreviations

SDMs: species distribution models
GARP: Genetic Algorithm for Ruleset Prediction
US: the United States
ENMs: ecological niche models
CART: classification and regression tree
DG: DesktopGARP
UI: Unimportance Index
AUC: area under the curve
BRTs: boosted regression trees

## References

Alexander, K.A., Lewis, B.L., Marathe, M., Eubank, S., Blackburn, J.K., 2012. Modeling of wildlife-associated zoonoses: applications and caveats. Vector-Borne Zoonotic Dis. 12, 1005–1018.

Anderson, R.P., Lew, D., Peterson, A.T., 2003. Evaluating predictive models of species’ distributions: criteria for selecting optimal models. Ecol. Model. 162, 211–232.

Araujo, M.B., Guisan, A., 2006. Five (or so) challenges for species distribution modelling. J. Biogeogr. 33, 1677–1688.

Austin, M., 2007. Species distribution models and ecological theory: a critical assessment and some possible new approaches. Ecol. Model. 200, 1–19.

Austin, M.P., Van Niel, K.P., 2011. Improving species distribution models for climate change studies: variable selection and scale. J. Biogeogr. 38, 1–8.

Barro, A.S., Fegan, M., Moloney, B., Porter, K., Muller, J., Warner, S., Blackburn, J.K., 2016. Redefining the Australian anthrax belt: Modeling the ecological niche and predicting the geographic distribution of Bacillus anthracis. PLoS Negl. Trop. Dis. 10, e0004689.

Blackburn, J.K., 2006. Evaluating the spatial ecology of anthrax in North America: Examining epidemiological components across multiple geographic scales using a GIS-based approach.

Blackburn, J.K., Asher, V., Stokke, S., Hunter, D.L., Alexander, K.A., 2014a. Dances with anthrax: wolves (Canis lupus) kill anthrax bacteremic plains bison (Bison bison bison) in southwestern Montana. J. Wildl. Dis. 50, 393–396.

Blackburn, J.K., Goodin, D.G., 2013. Differentiation of springtime vegetation indices associated with summer anthrax epizootics in west Texas, USA, deer. J. Wildl. Dis. 49, 699–703.

Blackburn, J.K., McNyset, K.M., Curtis, A., Hugh-Jones, M.E., 2007. Modeling the geographic distribution of Bacillus anthracis, the causative agent of anthrax disease, for the contiguous United States using predictive ecologic niche modeling. Am. J. Trop. Med. Hyg. 77, 1103–1110.

Blackburn, J.K., Van Ert, M., Mullins, J.C., Hadfield, T.L., Hugh-Jones, M.E., 2014b. The necrophagous fly anthrax transmission pathway: empirical and genetic evidence from wildlife epizootics. Vector-Borne Zoonotic Dis. 14, 576–583.

Elith, J., Leathwick, J.R., 2009. Species distribution models: ecological explanation and prediction across space and time. Annu. Rev. Ecol. Evol. Syst. 40, 677.

Fielding, A.H., Bell, J.F., 1997. A review of methods for the assessment of prediction errors in conservation presence/absence models. Environ. Conserv. 24, 38–49.

Friedman, J.H., Meulman, J.J., 2003. Multiple additive regression trees with application in epidemiology. Stat. Med. 22, 1365–1381.

Grinnell, J., 1917. The niche-relationships of the California Thrasher. The Auk 34, 427–433.

Hanley, J.A., McNeil, B.J., 1982. The meaning and use of the area under a receiver operating characteristic (ROC) curve. Radiology 143, 29–36.

Hay, S.I., Tatem, A.J., Graham, A.J., Goetz, S.J., Rogers, D.J., 2006. Global environmental data for mapping infectious disease distribution. Adv. Parasitol. 62, 37–77.

Hijmans, R.J., Cameron, S.E., Parra, J.L., Jones, P.G., Jarvis, A., 2005. Very high resolution interpolated climate surfaces for global land areas. Int. J. Climatol. 25, 1965–1978.

Hugh-Jones, M., Blackburn, J., 2009. The ecology of Bacillus anthracis. Mol. Aspects Med. 30, 356–367.

Hugh-Jones, M.E., De Vos, V., 2002. Anthrax and wildlife. Rev. Sci. Tech.-Off. Int. Epizoot. 21, 359–384.

Huston, M.A., 2002. Introductory essay: critical issues for improving predictions. Predict. Species Occur. Issues Accuracy Scale 7–21.

Hutchinson, G.E., 1957. Cold spring harbor symposium on quantitative biology. Concluding Remarks 22, 415–427.

Joyner, T.A., 2010. Ecological niche modeling of a zoonosis: A case study using anthrax outbreaks and climate change in Kazakhstan.

Kracalik, I.T., Kenu, E., Ayamdooh, E.N., Allegye-Cudjoe, E., Polkuu, P.N., Frimpong, J.A., Nyarko, K.M., Bower, W.A., Traxler, R., Blackburn, J.K., 2017. Modeling the environmental suitability of anthrax in Ghana and estimating populations at risk: Implications for vaccination and control. PLoS Negl. Trop. Dis. 11, e0005885.

Larson, S.R., Degroot, J.P., Bartholomay, L.C., Sugumaran, R., 2010. Ecological niche modeling of potential West Nile virus vector mosquito species in Iowa. J. Insect Sci. 10, 110.

Levine, R.S., Peterson, A.T., Yorita, K.L., Carroll, D., Damon, I.K., Reynolds, M.G., 2007. Ecological niche and geographic distribution of human monkeypox in Africa. PloS One 2, e176.

Levine, R.S., Yorita, K.L., Walsh, M.C., Reynolds, M.G., 2009. A method for statistically comparing spatial distribution maps. Int. J. Health Geogr. 8, 7.

Lim, B., Klein, K.J., 2006. Team mental models and team performance: A field study of the effects of team mental model similarity and accuracy. J. Organ. Behav. 27, 403–418.

Lobo, J.M., Jiménez-Valverde, A., Real, R., 2008. AUC: a misleading measure of the performance of predictive distribution models. Glob. Ecol. Biogeogr. 17, 145–151.

Martinez-Meyer, E., Peterson, A.T., Servín, J.I., Kiff, L.F., 2006. Ecological niche modelling and prioritizing areas for species reintroductions. Oryx 40, 411–418.

McNyset, K.M., Blackburn, J.K., 2006. Does GARP really fail miserably? A response to. Divers. Distrib. 12, 782–786.

Morris, L.R., Proffitt, K.M., Asher, V., Blackburn, J.K., 2016. Elk resource selection and implications for anthrax management in Montana. J. Wildl. Manag. 80, 235–244.

Mullins, J., Lukhnova, L., Aikimbayev, A., Pazilov, Y., Van Ert, M., Blackburn, J.K., 2011. Ecological Niche Modelling of the Bacillus anthracis A1. a sub-lineage in Kazakhstan. BMC Ecol. 11, 32.

Mullins, J.C., Garofolo, G., Van Ert, M., Fasanella, A., Lukhnova, L., Hugh-Jones, M.E., Blackburn, J.K., 2013. Ecological niche modeling of Bacillus anthracis on three continents: evidence for genetic-ecological divergence? PloS One 8, e72451.

Openshaw, S., Taylor, P., 1979. A million or so correlation coefficients: three experiments on the modifiable areal unit problem. 127–144. Stat. Appl. Spat. Sci. Pion Lond.

Ostfeld, R.S., Glass, G.E., Keesing, F., 2005. Spatial epidemiology: an emerging (or re-emerging) discipline. Trends Ecol. Evol. 20, 328–336.

Pearson, R.G., Dawson, T.P., 2003. Predicting the impacts of climate change on the distribution of species: are bioclimate envelope models useful? Glob. Ecol. Biogeogr. 12, 361–371.

Peterson, A., Cohoon, K., 1999. Sensitivity of distribution prediction algorithms to geographic completeness. Ecol. Model. 117, 159–164.

Peterson, A.T., Papeş, M., Eaton, M., 2007. Transferability and model evaluation in ecological niche modeling: a comparison of GARP and Maxent. Ecography 30, 550–560.

Peterson, A.T., Vieglais, D.A., 2001. Predicting Species Invasions Using Ecological Niche Modeling: New Approaches from Bioinformatics Attack a Pressing Problem: A new approach to ecological niche modeling, based on new tools drawn from biodiversity informatics, is applied to the challenge of predicting potential species’ invasions. BioScience 51, 363–371.

Pulliam, H.R., 1988. Sources, sinks, and population regulation. Am. Nat. 132, 652–661.

Sloyer, K., Burkett-Cadena, N.D., Yang, A., Corn, J.L., Vigil, S.L., McGregor, B.L., Wisely, S.M., Blackburn, J.K., 2018. Ecological niche modeling the potential geographic distribution of four Culicoides species of veterinary significance in Florida. bioRxiv 447003.

Stein, C.D., 1945. The history and distribution of anthrax in livestock in the United States. Vet Med 40, 340–349.

Stockwell, D., 1999. The GARP modelling system: problems and solutions to automated spatial prediction. Int. J. Geogr. Inf. Sci. 13, 143–158.

Sweeney, A., Beebe, N., Cooper, R., 2007. Analysis of environmental factors influencing the range of anopheline mosquitoes in northern Australia using a genetic algorithm and data mining methods. Ecol. Model. 203, 375–386.

Thomasson, V., Blouin-Demers, G., 2015. Using habitat suitability models considering biotic interactions to inform critical habitat delineation: An example with the eastern hog-nosed snake (Heterodon platirhinos) in Ontario, Canada. Can Wildl. Biol Manag 4, 1–17.

Van Ness, G., 1959. Anthrax—a soil borne disease. Soil Conserv 21, 206–208.

Van Ness, G.B., 1971. Ecology of anthrax. Science 172, 1303–1307.

Vega, G., Pertierra, L., Olalla-Tárraga, M., 2016. Data from: MERRAclim, a high-resolution global dataset of remotely sensed bioclimatic variables for ecological modelling. Dryad Digit. Repos.

Vega, G.C., Pertierra, L.R., Olalla-Tárraga, M.Á., 2017. MERRAclim, a high-resolution global dataset of remotely sensed bioclimatic variables for ecological modelling. Sci. Data 4, 170078.

